# Internal control for process monitoring of clinical metagenomic next-generation sequencing of urine samples

**DOI:** 10.1101/2020.03.27.012930

**Authors:** Victoria A. Janes, Jennifer S. van der Laan, Sébastien Matamoros, Daniel R. Mende, Menno D. de Jong, Constance Schultsz

## Abstract

**Background:** Process control for clinical metagenomic next-generation sequencing (mNGS) is not yet widely applied, while technical sources of bias are plentiful. We present an easy-to-use internal control (IC) method focussing on technical process control applied to metagenomics in clinical diagnostics.

**Methods:** DNA of nine urine samples was sequenced in the absence and presence of *Thermus thermophilus* DNA as IC in incremental concentrations (0.5-2-5%). Between aliquots of each sample, we compared the IC relative abundance (RA), and after *in silico* subtraction of IC reads, the microbiota and the RA of pathogens. The optimal IC spike-in concentration was defined as the lowest concentration still detectable in all samples.

**Results:** The RA of IC correlated linearly with the spiked IC concentration (r^2^=0.99). IC added in a concentration of 0.5% of total DNA concentration was detectable in all samples, regardless of human/bacterial composition and after *in silico* removal gave the smallest difference in RA of pathogens compared to the unspiked aliquot of the sample. The microbiota of sample aliquots sequenced in the presence and absence of IC was highly similar after *in silico* removal of IC reads (median BC-dissimilarity per sample of 0.059), provided samples had sufficient bacterial read counts.

**Conclusion:** *T. thermophilus* DNA at a percentage of 0.5% of the total DNA concentration can be applied for the process control of mNGS of urine samples. We demonstrated negligible alterations in sample microbial composition after *in silico* subtraction of IC sequence reads. This approach contributes toward implementation of mNGS in the clinical microbiology laboratory.

## INTRODUCTION

Metagenomic next-generation sequencing (mNGS) holds potential as a rapid pathogen detection tool for clinical microbiology diagnostics(1). In mNGS, all DNA in a sample has an equal chance of being sequenced, including DNA from host and contaminant cells. This means the interpretation of mNGS results can be challenging. For example, common reagent contaminants were previously incorrectly identified as causative of infection(2, 3). Differences in sample type, starting DNA concentration of a sample, library preparation efficiency for GC-rich organisms, and sequencing depth can all potentially bias the mNGS readout(4–7). If mNGS is to be safely used for clinical diagnostics, potential sources of error or variation in library preparation and sequencing need to be considered and controlled for.

While blank extraction controls and mock community and/or positive control samples can be used to assess the level of contamination and library preparation efficiency at batch level, external controls are not suitable for process monitoring of individual samples that vary greatly in microbial and host composition, DNA concentration and possible inhibitors. To this end, an internal control (IC) consisting of low concentration exogenous DNA that is completely unrelated to potential pathogens and the host microbiota, can be spiked into extracted DNA from each sample and subsequently detected nested within the diagnostic test procedure. Detection of a constant relative abundance (RA) of IC across all samples would ensure the library preparation and sequencing to be technically successful, ruling out technical error in samples where no pathogen reads are detected.

The application of ICs as process control is standard practise in molecular diagnostic microbiology such as for pathogen-specific PCR(8), but is still lacking in reported diagnostic applications of mNGS(2, 9–14). Studies that do describe ICs in metagenomic analyses typically applied fixed amounts of short synthetic DNA aimed at quantification of microorganisms or quantification of competition between target DNA and host DNA in library preparation protocols that include a DNA amplification step(2, 6, 15–18). It is unclear if the type and fragment length of IC used for such protocols are directly applicable to PCR-free sequencing protocols, such as the protocol used for diagnostic mNGS of clinical urine samples(19). Spike-in quantification standards are often patented and expensive while simpler and cheaper spike-ins may suffice for technical process control. Most importantly, because the DNA concentration of clinical urine samples varies greatly, spiking samples with a fixed amount of IC will result in overrepresentation of IC in low concentration samples and underrepresentation of IC reads in high concentration samples making such a strategy unsuitable for process control.

We describe the design and validation of an IC procedure for the process control of PCR-free library preparation and mNGS of clinical urine samples using full-length *Thermus thermophilus* DNA in a concentration that was titrated according to the DNA concentration of the urine sample.

## METHODS

### Urine sample collection

As part of a larger study of diagnostic mNGS of urinary tract infections (UTIs), we collected consecutive urine samples that were sent to the clinical microbiology laboratory of the Academic Medical Center, Amsterdam UMC for routine diagnostic semi-quantitative culture(19). Personal data were anonymised and handled in compliance with the General Data Protection Regulation and medical-ethical guidelines of the Academic Medical Center Amsterdam for anonymised use of patient materials. DNA extraction was performed after in-house enzymatic lysis, using the automated NucliSENS easyMag platform (Biomérieux) according to manufacturer’s instructions(19). These samples were sequenced without IC and the DNA was stored at −80°C until further use. Nine representative samples of this collection were selected based on variations in bacterial versus human read counts, culture results, and DNA concentration (table 1) and were sequenced again in the presence of incremental concentrations of IC. The selected samples represented the extremes, i.e. both culture positive and negative high human/low bacterial read samples (A, B, C), high bacterial/low human read samples (F, G), equal human/bacterial read samples (E), and culture negative samples with a low DNA concentration (H, I).

**Table 1.** Characteristics of the sequenced urine samples. The total number of reads, IC, unclassified, bacterial and human reads per sample aliquot are stated. Semi-quantitative culture results are given in colony forming units per ml of urine (CFU/ml).

| Sample | Culture result | DNA concentration (ng/μl) | % spiked IC DNA | Total no. of reads | No. of IC reads | No. of unclassified reads | No. of bacterial reads | No. of human reads |
| --- | --- | --- | --- | --- | --- | --- | --- | --- |
| A | *S. aureus* >10<sup>4</sup> CFU/ml | 12,8 | 0 | 1800197 | 0 | 54004 | 686 | 1745507 |
|  |  |  | 0.5 | 2834871 | 2598 | 3687 | 2761 | 2825825 |
|  |  |  | 2 | 2906634 | 13076 | 4006 | 11769 | 2877783 |
|  |  |  | 5 | 3476412 | 36171 | 7003 | 2124 | 3431114 |
| B | *P. aeruginosa* >10<sup>4</sup> CFU/ml, *E. cloacae* >10<sup>4</sup> CFU/ml | 17,68 | 0 | 1217911 | 0 | 31239 | 3697 | 1182975 |
|  |  |  | 0 | 4230673 | 42 | 5609 | 16060 | 4208962 |
|  |  |  | 0.5 | 3669184 | 6844 | 5283 | 13648 | 3643409 |
|  |  |  | 2 | 3146673 | 22956 | 5870 | 12104 | 3105743 |
|  |  |  | 5 | 3142635 | 58897 | 9333 | 11404 | 3063001 |
| C | Commensal flora 10<sup>4</sup> CFU/ml | 10,68 | 0 | 1222137 | 0 | 39799 | 16904 | 1165434 |
|  |  |  | 0.5 | 4060689 | 8404 | 10986 | 45521 | 3995778 |
|  |  |  | 5 | 3525448 | 62633 | 14092 | 38950 | 3409773 |
| D | Commensal flora >10<sup>4</sup> CFU/ml | 12,26 | 0 | 1018979 | 0 | 111370 | 94437 | 813172 |
|  |  |  | 2 | 2469179 | 32993 | 225690 | 244708 | 1965788 |
|  |  |  | 5 | 3107498 | 83231 | 271172 | 298285 | 2454810 |
| E | *E. coli* >10<sup>4</sup> CFU/ml | 11,68 | 0 | 1276427 | 2 | 45820 | 1222817 | 7788 |
|  |  |  | 0.5 | 3283575 | 8834 | 143238 | 3081986 | 49517 |
|  |  |  | 5 | 3891596 | 117764 | 176733 | 3534129 | 62970 |
| F | *E. coli* >10<sup>5</sup> CFU/ml, *K. pneumoniae* >10<sup>5</sup> CFU/ml | 22,00 | 0 | 1529193 | 0 | 57574 | 1423548 | 48071 |
|  |  |  | 0.5 | 3935399 | 3266 | 181040 | 3612043 | 139050 |
|  |  |  | 2 | 4386781 | 12872 | 201483 | 4013053 | 159373 |
|  |  |  | 5 | 3572208 | 29667 | 162892 | 3241806 | 137843 |
| G | *P. mirabilis* >10<sup>5</sup> CFU/ml, *M. morganii* >10<sup>5</sup> CFU/ml, *P. aeruginosa* <10<sup>4</sup> CFU/ml | 9,78 | 0 | 1427110 | 0 | 56789 | 649441 | 720880 |
|  |  |  | 5 | 2860345 | 54974 | 71919 | 1285320 | 1448132 |
| H | Commensal flora 10<sup>4</sup> CFU/ml | 8,2 | 0 | 1395035 | 0 | 17319 | 15325 | 1362391 |
|  |  |  | 5 | 3628947 | 76747 | 29465 | 96711 | 3426024 |
| I | Commensal flora 10<sup>4</sup> CFU/ml | 7,94 | 0 | 1624302 | 0 | 108709 | 12326 | 1503267 |
|  |  |  | 2 | 2780217 | 29602 | 117812 | 23174 | 2609629 |
| IC | *T. thermophilus* | 1,00 | 100 | 2680871 | 2425350 | 199119 | 10558 | 45844 |

### IC procedure

We designed an IC with comparable GC-content and genome length to the common Gram-negative uropathogen *Escherichia coli* (~5.1Mb; GC% 50.6; NCBI:taxid 562). Further, the IC should not be a human pathogen or part of the human microbiota or have genomic similarity to these. The full-length bacterial genome of the extremophile *Thermus thermophilus* met these criteria and was previously successfully used as IC in analyses of environmental river microbiota(20).

To allow for detection of IC DNA at a low constant RA across all samples, we titrated the IC concentration relative to the sample DNA concentration. Therefore, DNA was first extracted from urine and quantified before addition of IC DNA at an appropriate concentration relative to the total DNA concentration.

DNA concentrations of stored DNA aliquots (−80 C) of each sample were measured using the Qubit HS dsDNA quantification kit (Thermo Fisher Scientific). The IC DNA concentrations were titrated to a final concentration of 0.5%, 2%, and 5% of the total DNA concentration of the sample. Not all concentrations were tested in each sample if limited amounts of DNA were available. IC DNA and sample DNA were mixed to a final DNA concentration of 1 ng/μl in a volume of 100μl (supplementary table 1). To assess variation between sequencing runs, two aliquots of the same sample were sequenced without IC in both runs (B and B1 + 0% IC sample, Table 1). In addition, B1+0% IC sample enabled us to see which proportion of reads were derived through contamination as this was the only sample sequenced without IC on the 2^nd^ run. A sample only containing IC (*T. thermophilus* DNA suspended in TE buffer 1x, 1ng/μl) was sequenced to confirm that *T. thermophilus* reads were not misidentified as other taxa.

### Library preparation and sequencing

Sample aliquots were sheared into 200bp fragments using the S220 Focused-ultrasonicator (Covaris). The Ion Xpress™ Plus Fragment Library Kit (Thermo Fisher Science) was used for PCR-free, library preparation according to manufacturer’s specifications. Sequencing was performed on the Ion Torrent Proton and PGM (Thermo Fisher Scientific). This platform was used because of its sequencing speed which is required for clinical diagnostics. The platform was set to produce 2 million reads per sample for run 1 (10 samples sequenced as part of 55 samples in total) and 3 million reads per sample for run 2, both 200bp single-end (SE).

### Sequence analysis

We used FastQC to check the read quality(21), Trimmomatic V0.38 to remove low quality reads(22), and BBMap to quantify and remove human reads by aligning to the GRCh38 Human Genome obtained from Genbank(23, 24). The IC and bacterial reads were identified using Kraken2 against the MiniKraken2_v1_8GB database, downloaded on the 2^nd^ July 2019(25). The RA of IC was defined as the *Thermus* read count at genus level divided by the total number of reads per sample (including human and unclassified reads). We assessed the RA of the IC per sample and per spiked IC DNA concentration to identify the lowest IC DNA concentration that was still detectable across all samples. We assessed the RA of IC in the B1+0% sample to assess the proportion of IC reads derived through cross-contamination. Next, all reads mapping to the genus *Thermus* were removed *in silico* before we used Bracken to calculate the RA of all classified bacterial species by Bayesian re-estimation of sequence reads(26). Bacterial species were annotated as urinary pathogens based on clinical microbiology common practice as well as the results of a scoping review of the literature (Supplementary table 2).

For each sample, the RA of urinary pathogens was compared between aliquots sequenced in the presence and absence of IC. In case of a polymicrobial composition, the cumulative RA of pathogens was used. The cumulative RA of pathogens was compared between sample aliquots sequenced in the presence and absence of IC after *in silico* subtraction of IC reads. To assess whether sequencing samples in the presence of IC and subsequent *in silico* subtraction of IC reads affected the sample microbiota, the RA of each detected species was compared between aliquots by calculating the Bray-Curtis dissimilarity (BC-dissimilarity). The optimal spike-in concentration of IC DNA was defined as the lowest still detectible IC concentration with minimum impact on the microbiota and cumulative RA of pathogens.

## RESULTS

### Sequencing results

We sequenced 28 aliquots of DNA obtained from 9 urine samples, spiked with incremental IC concentrations, in addition to one sample containing 100% IC DNA. The mean read count per sample aliquot was 1,486,600 (SD=252,004) for the unspiked samples of the first run, and 4,093,176 reads (SD = 615,999) for the spiked samples sequenced in the second run (table 1). The read distributions per sample are depicted in figure 1. In the samples with high human read count (samples A-D, H and I), the mean human read count varied between 1,2-1,4 million reads and the mean bacterial read count varied between 2,442-244,708 per sample. For the samples with high bacterial read count (samples E and F) the mean human read count per sample was 44,517 and 138,446 versus 3 and 3,4 million bacterial reads respectively. Resequencing the sample with equal bacterial/human read count (sample G) again produced an equal distribution of human and bacterial reads (figure 1). Two aliquots of sample B were sequenced without IC on both runs. The RA of bacterial species of the two aliquots correlated strongly (BC-dissimilarity 0.061), indicating results from the two sequencing runs were comparable.

**Figure 1.**
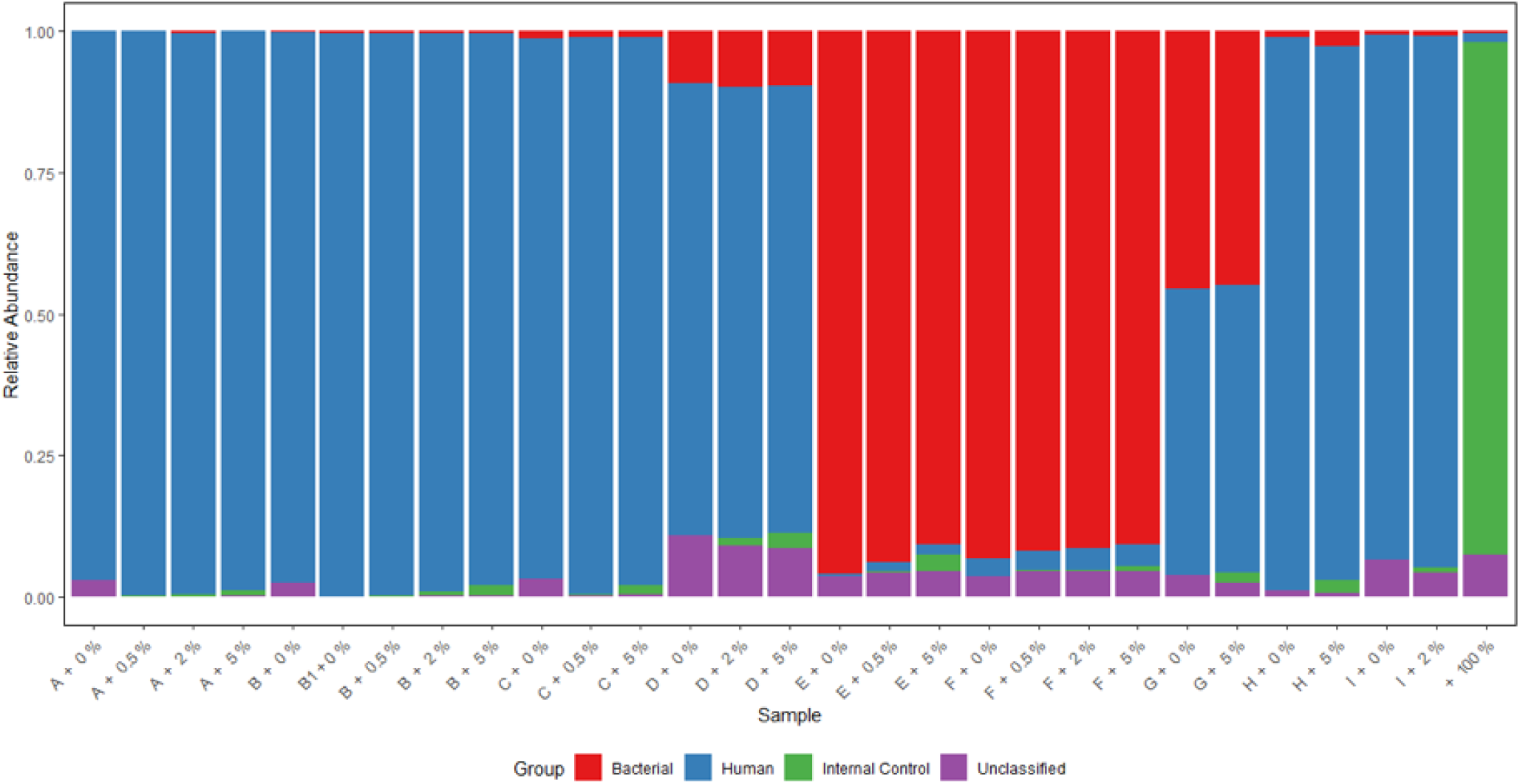
Read distributions. Shown is the total read distribution of each sequenced aliquot, indicating the RA of bacterial (red), human (blue), IC (green) and unclassified (purple) reads. Following the sample name (A – IC) is the spiked IC concentration (%).

### Detection of IC

The resequencing of an aliquot of sample B without IC on the 2^nd^ run (amidst samples sequenced in the presence of IC) not only allowed assessment of between-run variability but was also used to assess the proportion of IC reads derived through contamination. Out of 4,230,673 reads in this aliquot (aliquot name B1+0%), 42 reads (0.001%) identified as *Thermus* at genus level of which 39 identified as *T. thermophilus*.

To assess whether *T. thermophilus* reads mapped against other bacterial species, a sample containing 100% IC DNA was sequenced. Of all 2,635,027 reads, 2,425,350 reads (92%) mapped to the *Thermus* genus and of these 1,885,710 (71,6% of total) were classified as *T. thermophilus*. 2,544 (0.096%) reads mapped to other bacterial species, predominantly to *E. coli* and *Klebsiella pneumoniae*, 8,014 reads (0.3%) mapped to the domain Bacteria, 45,844 reads identified as human (1.7%), and 199,119 (7,5%) reads remained unclassified.

Next, we assessed the RA of IC reads per sample at IC DNA concentrations of 0.5, 2 and 5% of the total DNA concentration. The RA of IC reads rose linearly with increasing concentrations of spiked IC DNA (Pearson’s r^2^=0.99) and was 10.24 times higher in the samples spiked with IC DNA at 5% of the total DNA concentration compared to the samples spiked with IC DNA at 0.5% of the total DNA concentration (figure 2). There was no significant difference in RA of IC between samples with a high or equal bacterial read count (E, F and G) compared to samples with a low bacterial and high human read count (A-D, H and I) (Welch two sample t-test, p = 0.8).

**Figure 2.**
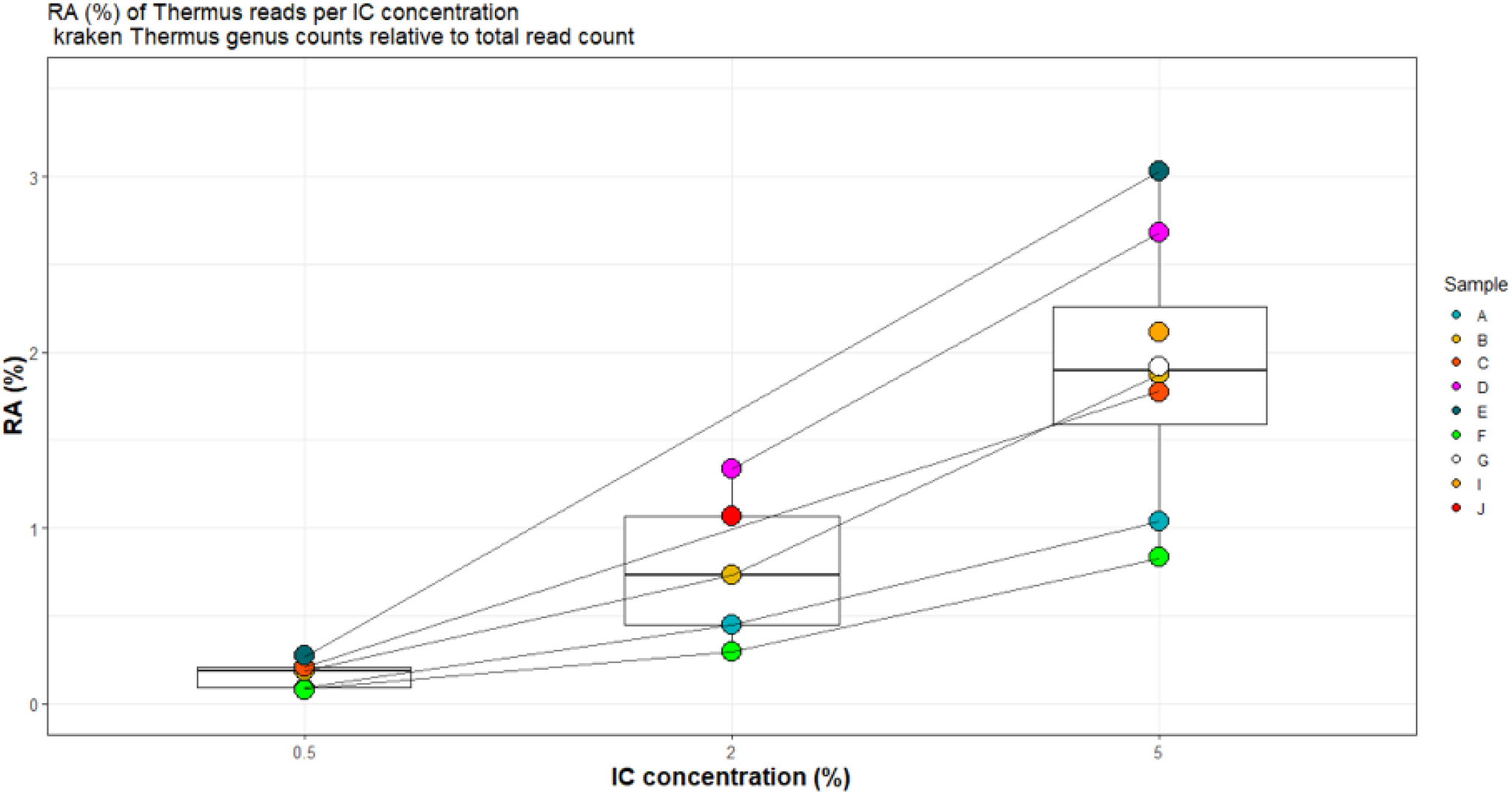
**RA of IC per sample and per IC concentration.**

IC DNA spiked into samples at concentration of 0.5% of the total DNA concentration was detectable in all samples (median RA 0.19%; IQR 0.12%; range 0.09-0.27%). The median RA of IC reads at this concentration was 187 times higher than the RA of the IC reads in the aliquot of sample B that was sequenced in the absence of IC on run 2, and thus clearly exceeded the RA of IC expected to be derived through cross-contamination.

After *in silico* subtraction of IC reads, the RA of the bacterial species reads was highly similar between sample aliquots sequenced in the presence and absence of IC, illustrated by the similar species distribution between aliquots of the same sample seen in figure 3 and by the median BC-dissimilarity per sample of 0.059 (IQR 0.04, range 0.008-0.61). The exception was sample A with a BC-dissimilarity of 0.61. The aliquots of this sample had a median bacterial read count of only 2442 (range 686-11,769), over 61 times lower than the median bacterial read count of all other samples (median 150,363; range 12,043 (sample B) – 3,426,924 (sample F)).

**Figure 3.**
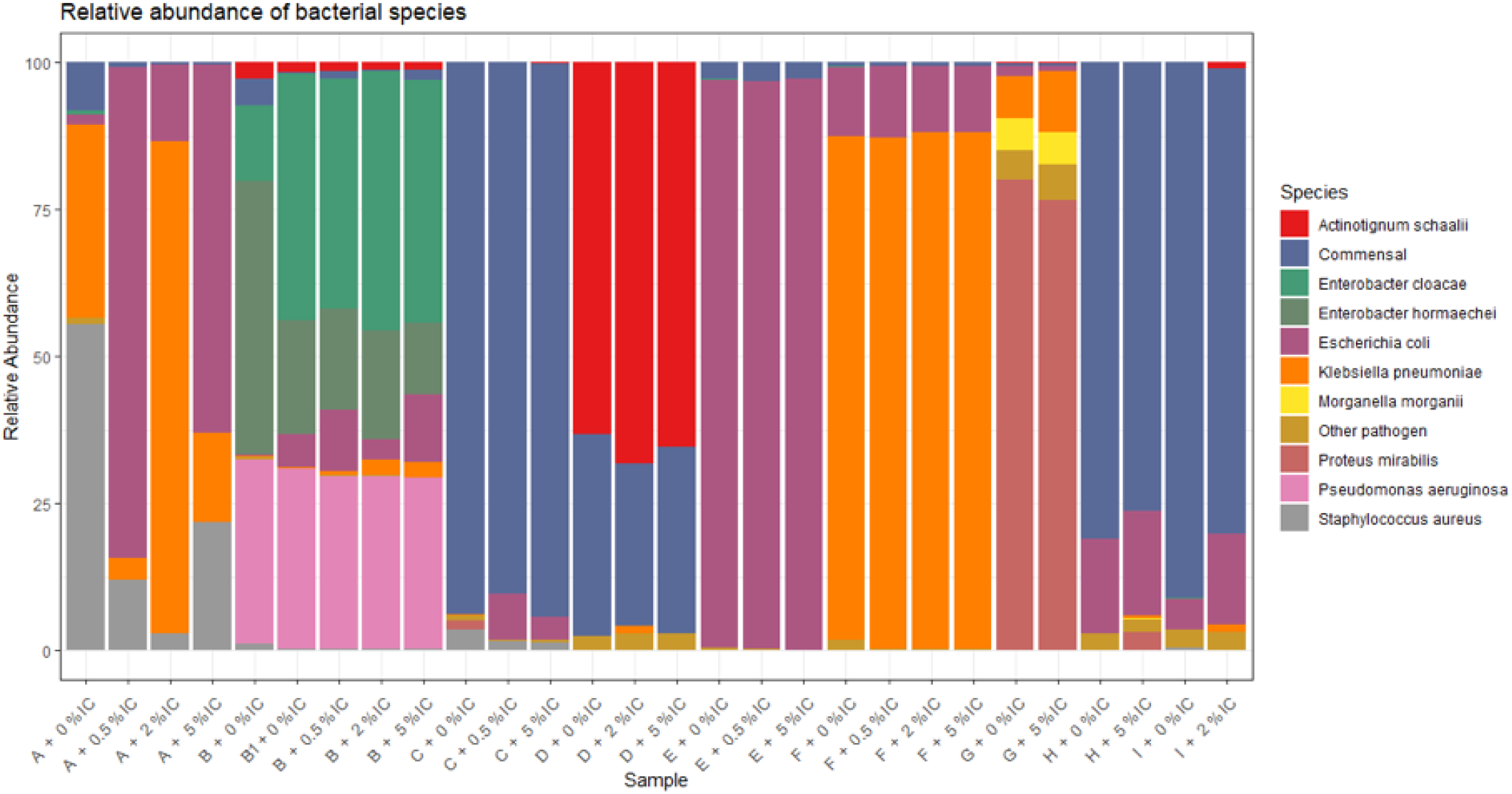
Relative abundance of bacterial pathogenic species and commensals reads per sample aliquot. Shown is the RA of the 9 most abundant bacterial pathogens classified after *in silico* removal of IC. Commensal bacterial species were grouped (ochre yellow).

Since the main outcome measure for diagnostic mNGS of urine samples was the RA of bacterial pathogens, we calculated the difference in RA of bacterial pathogens between spiked and unspiked sample aliquots after *in silico* subtraction of IC reads (table 2). An IC DNA concentration of 0.5% gave the smallest difference in cumulative RA of bacterial pathogens between spiked and unspiked sample aliquots, and was still detected in all samples. The median difference in RA was 1.15% (table 2).

**Table 2.**
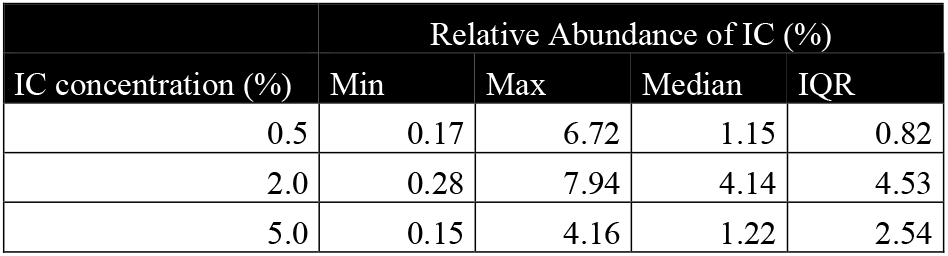
The median difference in cumulative RA of pathogens between spiked and unspiked aliquots of a sample after *in silico* removal of IC reads. Results are shown per IC concentration.

## DISCUSSION

In this study we successfully applied *T. thermophilus* DNA as IC for the process control of mNGS of clinical urine samples. DNA aliquots extracted from urine samples were sequenced in the absence and presence of incremental concentrations of IC DNA. IC DNA added at a concentration of 0.5% of the total sample DNA concentration was detectable in all samples, did not alter the microbial composition of the sample DNA or the RA of detected bacterial pathogens substantially after *in silico* removal of IC reads. The results presented here are directly relevant to clinical microbiology practise as they are derived from clinical urine samples, representing the full array of variation seen in microbial and host DNA composition and concentration.

Other studies described the use of fixed amounts of short fragment synthetic DNA as IC aimed at quantification or as a measure for pathogen detection sensitivity(2, 6, 15–17). For such uses of IC, the RA of IC in the readout is integral to the analysis, making this a different approach to using IC as process control, where it is desirable to detect a constant low RA of IC in each sample to ensure the sequencing process was technically successful. The DNA concentration of urine samples is highly variable, meaning spiking a fixed amount of IC, as done in these other studies, can easily lead to over- or underrepresentation of IC depending on the sample DNA concentration. Moreover, short fragments of IC could be sheared to even shorter fragments during library preparation and could consequently be lost during size selection when targeting species that require longer fragmentation time than the IC(20). The spiking of samples with IC in a concentration relative to the total DNA concentration of that sample, ensured detection of IC regardless of the concentration of the sample and thus allowed for consistent quality control of each sample.

Many bacteria that cause UTI, such as *E. coli*, are commensal to the genito-urinary tract and are only considered causative of disease when present in concentrations above a certain threshold(27). This study was not designed to address quantification, but quantification of urine mNGS results is needed to align with current routine culture-based microbiology diagnostics whereby a diagnosis is established based on semi-quantitative cultures.

Two reads were initially identified as *T. thermophilus* in sample D+0% sequenced on run 1 in the absence of IC. No samples were spiked with IC on this run. We hypothesised that this finding was caused by misclassification of Taq-polymerase DNA, derived from *Thermus aquaticus*, which may have regions of genomic similarity to *T. thermophilus*. However, BLASTn identified these 2 reads as *E. coli* plasmids p94EC-5.

When assessing the clinical relevance of species present at very low relative abundance, the likelihood that these reads are present as a result of cross-contamination should be taken into consideration. Relative abundances of 0.05-2.78% have been described for droplet cross-contamination during sample preparation(28). Barcode hopping, the incorrect assignment of library molecules from the expected barcode to a different barcode in a multiplexed pool, was observed in biological mock community samples at a rate of 0.033% on the IonTorrent PGM platform which has similar technology to the IonTorrent Proton(29). We demonstrate that the inclusion of IC DNA allowed for the assessment of cross-contamination by sequencing replicates in the presence and absence of IC. The impact of cross-contamination and barcode hopping appeared minimal in our setting. We propose that such a sample sequenced in the absence of IC should be included in each library preparation and sequencing run, to assess cross-contamination and to determine the minimum threshold of the RA of IC to consider a sample successfully sequenced.

By resequencing biological replicates in the presence and absence of IC divided over 2 sequencing runs, we demonstrated negligible alterations in sample microbial and pathogen composition after *in silico* subtraction of IC reads for 8 out of 9 samples. The incongruent sample A consisted of >95% human reads, leaving only a median of 2,442 (range 686-11,769) classified bacterial reads per sequenced aliquot. The low bacterial read count can explain the poor correlation between sequenced aliquots and this data may help set guidelines for a minimum sequencing depth for clinical samples and stresses the need for removal of host cells prior to sequencing(28, 29). Despite the variation in RA of bacterial species in sample A, the IC was detected in all aliquots, indicating sequencing and library preparation processes had been technically successful.

Our study has some limitations. Not every sample could be sequenced with each IC concentration, because limited volumes of DNA were present for some samples. However the overall results were consistent and RA of IC correlated linearly with spiked IC concentrations, implying our results can be extrapolated to other concentrations. Even lower spike-in concentrations could be suitable but were not tested in sufficient numbers. One aliquot of sample B was spiked with 0.05% IC DNA, yielding a RA of 0.025% (747 reads; data not shown).

Our approach does not control for DNA extraction efficiency. To achieve this, samples could be spiked with bacterial cells prior to DNA extraction. However, this comes with additional pitfalls. Spiking samples with a quantified bacterial cell suspension in a concentration of 0.5% relative to the sample will not necessarily result in 0.5% IC DNA. The human genome is several orders of magnitude longer than bacterial genomes, meaning each human cell produces approximately 1000 times more DNA than each bacterial cell(6). Given the number of sequencing reads produced per sequencing run is finite, host DNA will easily outcompete bacterial DNA, leading to difficulty in interpreting the mNGS read out: in cases where no pathogen reads are detected, distinguishing between a true negative (at the given sequencing depth) and a technical fail would be impossible. The spiking of samples with IC DNA in a concentration relative to the total DNA concentration of that sample mitigated this problem and ensured detection of IC regardless of the concentration or composition of the sample and thus allowed for consistent process control of each sample.

In conclusion we developed an IC for the process control of mNGS of urine samples that at a percentage of 0.5% of the total DNA concentration was detectable in all samples, regardless of the sample composition or DNA concentration. By resequencing sample aliquots in the presence and absence of IC, we demonstrated negligible alterations in sample microbial and pathogen composition after *in silico* subtraction IC. We showed how a lower detection threshold for detection of IC can be established taking contamination into account. This approach could be used as process control for diagnostic mNGS and contributes toward implementation of mNGS in the clinical microbiology laboratory.

## Acknowledgements

This study was supported by the COMPARE Consortium, which received funding from the European Union’s Horizon 2020 research and innovation programme under grant agreement No. 643476. The authors declare no conflict of interest.

